# Effectiveness of micromorphy against drilling predation: Insights from early Miocene faunal assemblage of Quilon limestone, India

**DOI:** 10.1101/856260

**Authors:** Debarati Chattopadhyay, K. S. Venu gopal, Devapriya Chattopadhyay

## Abstract

The nature of drilling predation, although well documented for molluscan fossils, is understudied for micromolluscs (<5mm). Studying predation in micromolluscs is especially critical in evaluating the adaptive significance of micromorphy against predation and assessing the importance of predator-prey size relationship (PPSR). This study documents drilling predation event in microbivalves from early Miocene (Burdigalian) fossil assemblage of Quilon limestone from Kerala, India. Our sample of ∼2000 valves represent nine families with an average drilling frequency (DF) of 0.06 and an incomplete drilling frequency (IDF) of 0.26. The characteristic drillhole morphology and occurrence of five genera of modern drilling gastropods (Naticid: *Natica, Tanea* and *Polinices*; Muricid: *Triplex* and *Dermomurex*) from the same locality reveals the predator identity. Predation in the studied assemblage is found to be highly selective in terms of prey taxa, size, mobility and site selection. Six out of nine families show evidence of predation indicating taxon selectivity. Poor correlation between DF and abundance further supports this view. Failed attacks are strongly correlated with morphological features such as surface ornamentation (Lucinidae), presence of conchiolin layers (Corbulidae). Drilling occurs primarily on medium size class and prey outside this size range show lower rate of attack. This indicates the existence of an “inverse size refugia” for extremely small prey along with the classical size refugia existing for large prey. Mobility is found to be a deterrent to drilling predation and it also increases failure.

Microbenthos of Quilon limestone shows a lower predation intensity in comparison to the Miocene macrobenthos worldwide including coeval formation of the Kutch Basin. The interaction in microbenthos is more strongly size-dependent in contrast to the Kutch fauna. Reduced predation intensity in microfauna and existence of “inverse size refugia” support the claim of micromorphy acting as a defense mechanism and highlights the role of size-dependent predation in marine benthos.

## Introduction

Predation is considered as one of the primary ecological processes that drives natural selection [1–3]. Predator-prey interactions are inherently dependent on size [4]. Consequently, body size of interacting predator and their prey are one of the vital predictors in controlling the feeding relationship within food web [5–7]. Based on global data on body size, Brose et al (2006) suggested that predator-prey size relationships (PPSRs) may systematically differ among habitats, such as terrestrial and marine [8]. The nature of PPSRs of marine invertebrates in deep time is especially interesting because of its impact on major evolutionary breakthrough driven by predation [1].

Drilling predation is one of the unique scenarios where the signature of the predatory event is preserved and various aspects of predator-prey dynamics can be reconstructed by evaluating non-random attacks based on species identity, ecological character, and size of the prey (see [9] for review). Such records are found in the recent [10–13] as well as in the fossil ecosystems dating as far back as Cambrian. The predation intensity varied during Phanerozoic, often in sync with diversity [14] and shows a significant increase from Late Mesozoic with the appearance of two modern predatory gastropod families, namely Muricid and Naticid [9, 15, 16].

It has long been recognized that predators are generally larger than their prey [17, but see 18], especially in highly size-structured marine ecosystem [19–21]. Size selectivity is largely true for drilling predator-prey system [22–24] with some exceptions [25]. The prey selection by a drilling predator is a balance between invested energy (through foraging, drilling and consumption) and energy gain (dependent on prey size) [22, 26]. As predator size increases, metabolic demand becomes more resulting in an increase in the rate of food intake to match the energetic demands [27]. Such increase in energy requirement can be tackled by choosing either a larger prey or attacking multiple smaller preys; the decision depends on the availability and distribution of prey-size in a community [28]. Prey often develops anti-predatory strategies as a response to an increase in predation pressure. Increase in effective size and attaining a “size refugia” is one of the common anti-predatory strategies exploited by marine invertebrates [29–33]. Such “size refugia” is often a result of handling limit of the predator [22]. Interestingly, prey smaller than the “size refugia” are not always attacked with equal frequency. It has been observed that the medium size class are often attacked the most, making them the preferred size class for predation [13, 25]. This points to an apparent predation-resistance of extremely small sized prey. If the attacks on extremely small prey significantly reduces the net energy gain of the predator, it would be expected to find a low predation pressure in smaller sized prey [27]. This may lead to an “inverse size refugia” in the smaller size class and shield the extremely small prey from predation. Harper and Peck [34] also demonstrated low intensity of durophagous predation in tropical brachiopods and attributed the micromorphic nature of tropical brachiopods as a defense against durophagous predation. Unfortunately, the drilling predation dynamics in juveniles and extremely small molluscan prey are rarely studied and hence, not well understood to test the existence of predation resistance in smaller prey.

The Quilon limestone of early Miocene, exposed in southern state of Kerala, India preserves a molluscan assemblage, dominated by bivalves and gastropods. This fauna is characterized by extremely small size (<5mm) in comparison to the bivalve fauna of coeval Chhasra formation of the Kutch Basin, India representing the same biogeographic province namely Western Indian Province (WIP) [35]. Both of these fauna displays abundance of predatory drillholes. Using these two fauna, we attempted to evaluate predator-prey dynamics in extremely small size class, addressing the following questions:

i. What is the nature of prey-predator dynamics (in terms of prey selectivity, size selectivity, site selectivity) in the extreme size class?
ii. Does the micromorphy provide any adaptive advantage against drilling predation?

## Materials and methods

### Geological setting and collection

All the specimens are collected from an extended cliff section of Asthamudi Lake (N 08°58’36”, E 076°38’08”) near Padapakkara village, Kerala, India (Fig 1). It corresponds to the Quilon limestone outcrop studied by Dey [36], Menon [37, 38], and Reuter et al [39]. This unit is represented by fossiliferous hard greenish limestone and interpreted as a seagrass habitat of Burdigalian (early Miocene) age [39]. The diverse fauna shows exquisite preservation of various invertebrate groups (including gastropod, bivalves, cephalopod, scaphopod, ostracodes, foraminifera and crabs) and dominated by individuals of small body size (<5 mm). Bulk sample was collected from the vertical face of the exposure.

**Fig 1.**
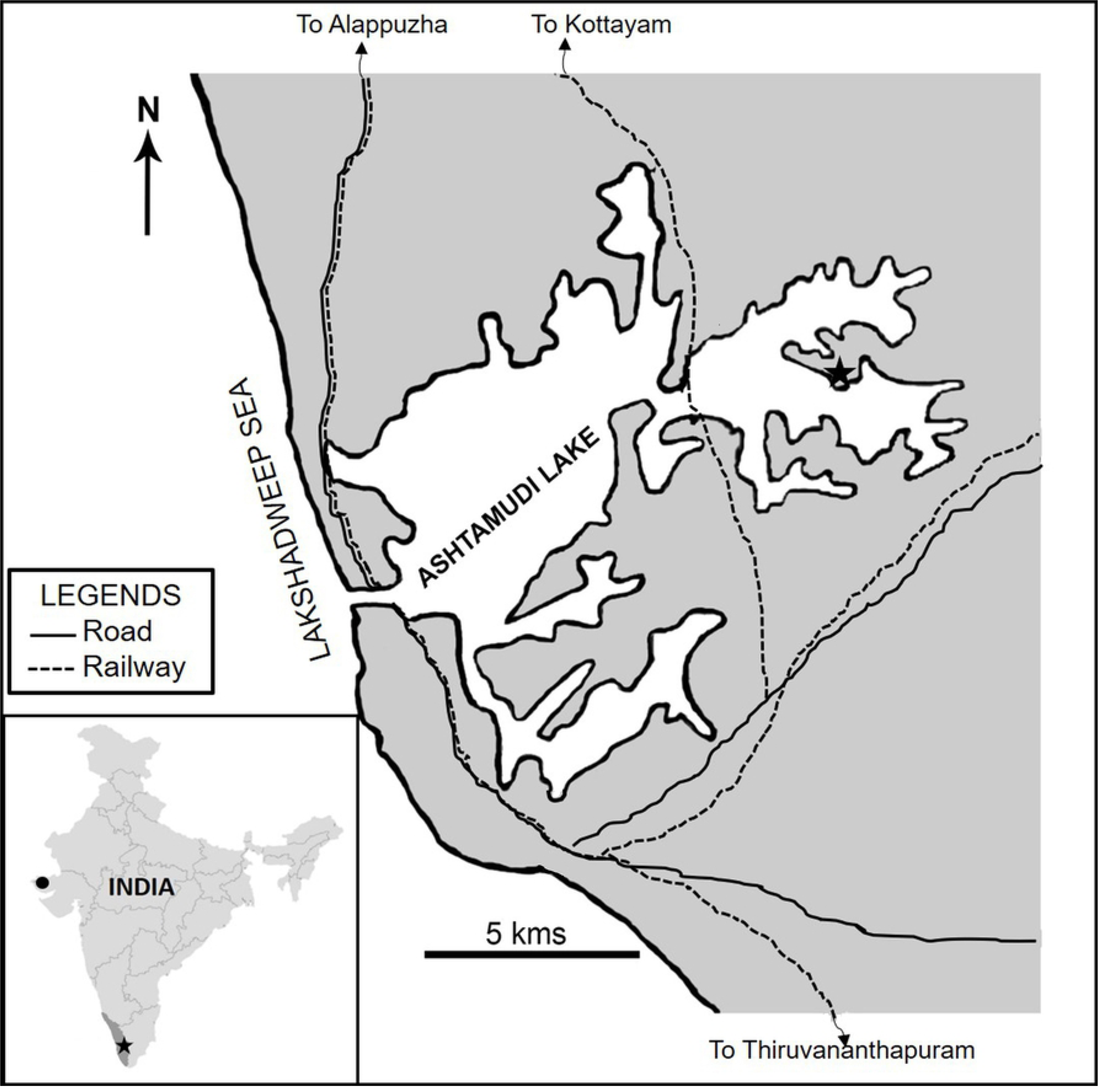
Map of Indian subcontinent with the Kerela and the Kutch Basin demarcated by a star and a circle respectively (inset). The detailed map of studied locality with exposures of Quilon limestone. The locality of collection is marked by a star.

Bulk sample of loose sediments (∼230gm) were soaked for 2-3 days in normal water. Repeated heating and thawing was used to separate specimens from the limestone matrix. The processed samples were then sieved using a stacked sieve of five mesh sizes (63, 60, 35, 25, 18μm). Sediments below 25μm sieve size did not contain any fossil. A total of 2032 intact bivalve specimens were studied under microscope and identified upto family level. Using the digitized images, the specimens were measured with the help of Image J. We measured all drilled specimens and 20 random undrilled specimens from each of the family for size analysis (Data File S1). We used SEM image to document representative specimens. We also compared our data with previously published predation data from the bivalve fauna of coeval formations including Chhasra formation of Kutch, India [25] (Fig 1).

### Analysis

Using the standard criteria for identifying predatory drillhole [16], we identified the predatory drillholes. We also found a few very small, cylindrical drillholes. The morphology of these drillholes is significantly different from the predatory drilling and hence, excluded from our analysis. We considered families that are represented by more than ten individuals for this study.

Because all the bivalve specimens in our collection are disarticulated valves, drilling frequency (DF) was calculated by dividing the number of drilled valves by the half of total number of valves in the collection [40].

Drilling frequency (DF) = N_D_ / (N ∗ 0.5)

Where

N_D_ = number of valves with complete drillhole

N = total number of valves.

The incomplete drilling frequency (IDF) was calculated by dividing the total number of incompletely drilled valves by the total number of drilled valves (complete and incomplete) present in the collection [25].

Incomplete drilling frequency (IDF) = N_ID_ / (N_ID_ + N_D_)

Where

N_ID_ = number of valves with incomplete drillhole

N_D_ = number of valves with complete drillhole.

Because there is no incidence of multiple drillholes for single specimen in our specimens, the calculated IDF is comparable to prey effectiveness (PE) proposed by Vermeij [1].

To reconstruct the predator size from a drill hole, we used following formulas proposed for Naticid [22] and Muricid [24] gastropods.

For Naticid, log (Y_nat_)= −0.372 + 0.552 (log X_nat_)

Where

Y_nat_= Drillhole diameter

X_nat_= Predator size

For Muricid, log_e_ (Y_mur_) = 0.82 log_e_ (X_mur_) – 2.46

Where,

Y_mur_=Maximum outer diameter of the drillhole

X_mur_=Maximum predator size.

We found it difficult to follow the standard protocols using sector grids [41] to assign location of the drillhole in the small specimens of the studied fauna. Instead, we divided each valve into three concentric regions (umbonal, middle and edge) and assigned the locations to the drillholes. The division of the sites was maintained consistently for all the drilled specimens.

We used Mann-Whitney U-test to compare response (in terms of size, drilling frequency) between various groups based on species identity, ecological character. To compare between different ecological groups and different size classes (small, medium, large) we used Chi^2^ test. To compare the size distribution of various categories (i.e. drilled, undrilled), we used Kolmogorov-Smirnov (K-S) test. To assess the relationship between predator and prey size, we used Pearson correlation test. All the analysis is done on R software [42].

## Results

### Predation intensity and success

A total of 2032 valves represent nine families of bivalves, dominated by Cardiidae (Fig 2, Fig 3A). A total of 62 valves show drillholes representing six families; Arcidae, Veneridae and Tellinidae does not have any drilled individuals (Table 1). The pooled DF is 0.06 and IDF is 0.25. We did not find incidence of multiple drillhole in any specimen. The majority of the drillholes are created by Naticid gastropods (84%).

**Fig 2.**
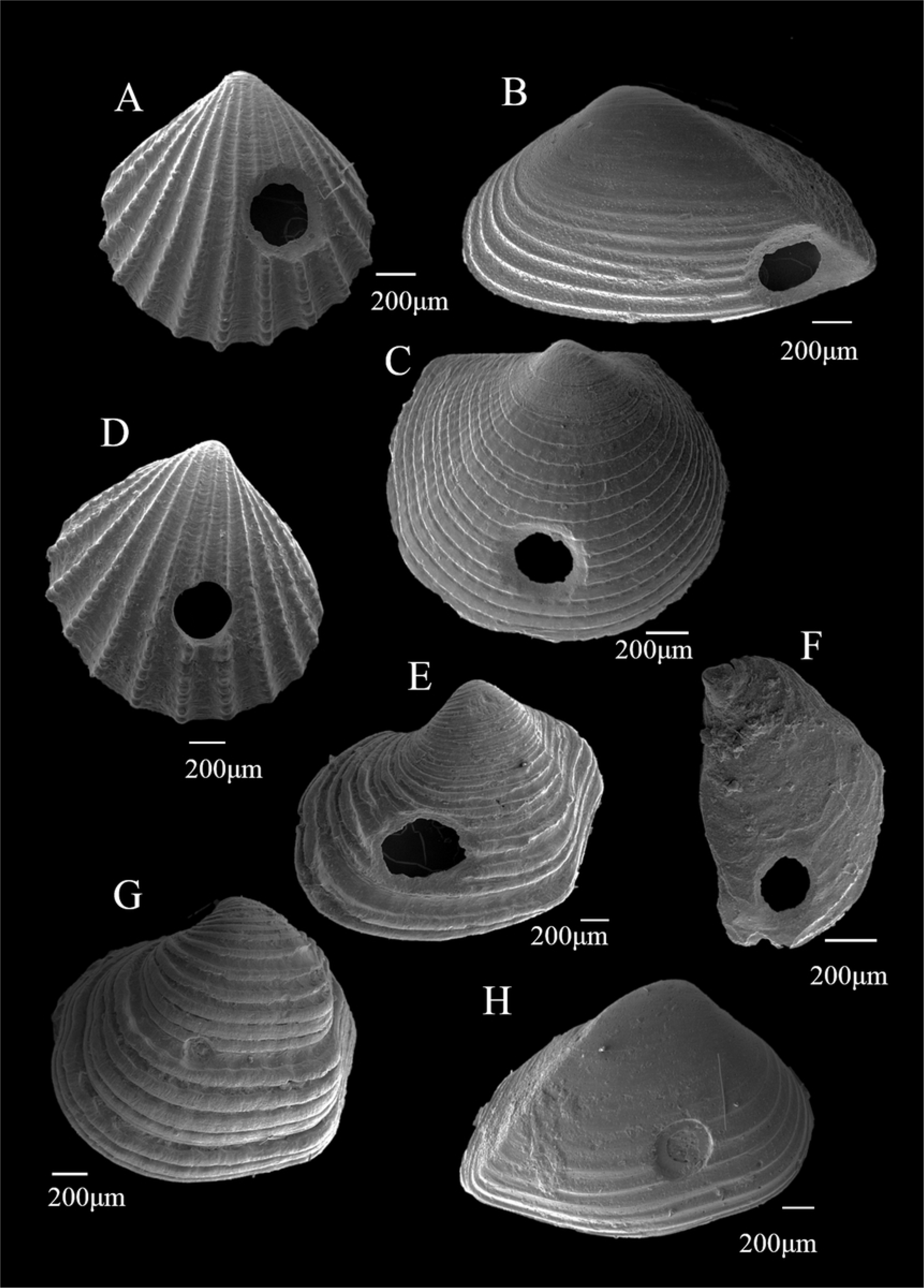
SEM pictures of the drilled bivalve families of Kerala, India. Complete drillholes are present in Cardiidae (A), Corbulidae (B), Glycymerididae (C), Cardiidae (D), Lucinidae (E), Anomidae (F) and incomplete drillhole on Lucinidae (G), Corbulidae (H).

**Fig 3.**
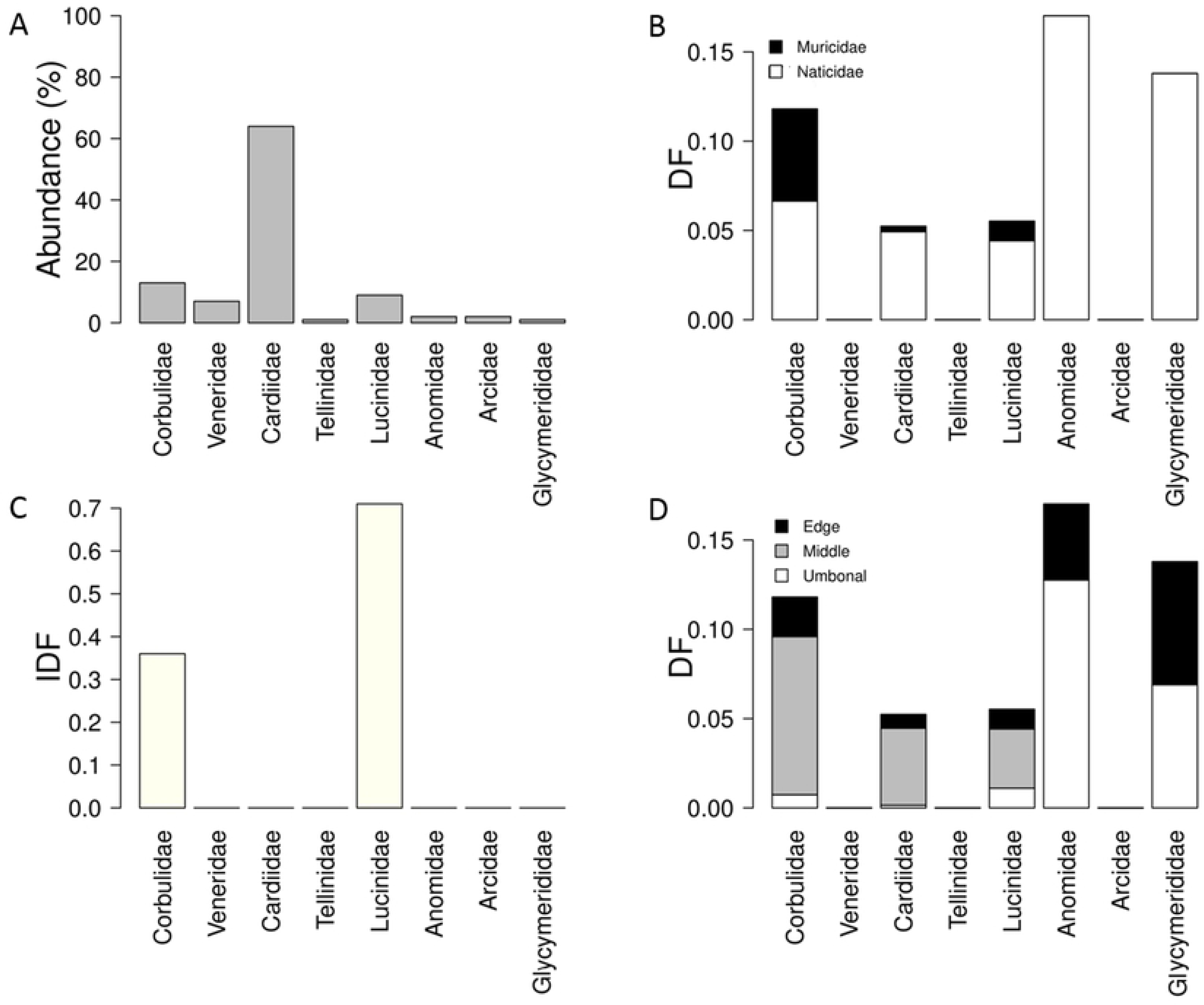
Plots showing families in the studied assemblage with their individual abundance (A); drilling frequency (DF) (B); incomplete drilling frequency (IDF) (C) and site selectivity (D).

**Table 1.** Taxonomic summary of drill hole data with ecological information of Kerala, India.

| Family | Abundance | Undrilled | Complete Drill holes | Drill holes at umbo | Drill holes at middle | Drill holes at edge | Naticid | Muricid | Incomplete Drill holes | DF | IDF | Substrate | Mobility |
| --- | --- | --- | --- | --- | --- | --- | --- | --- | --- | --- | --- | --- | --- |
| Corbulidae | 271 | 246 | 16 | 1 | 12 | 3 | 9 | 7 | 9 | 0.12 | 0.36 | Infaunal | Immobile |
| Veneridae | 147 | 147 | 0 | 0 | 0 | 0 | 0 | 0 | 0 | 0 | - | Infaunal | Mobile |
| Cardiidae | 1298 | 1264 | 34 | 1 | 28 | 5 | 32 | 2 | 0 | 0.05 | 0 | Infaunal | Mobile |
| Tellinidae | 23 | 23 | 0 | 0 | 0 | 0 | 0 | 0 | 0 | 0 | - | Infaunal | Mobile |
| Lucinidae | 181 | 164 | 5 | 1 | 3 | 1 | 4 | 1 | 12 | 0.06 | 0.71 | Infaunal | Mobile |
| Anomidae | 47 | 43 | 4 | 3 | 0 | 1 | 4 | 0 | 0 | 0.17 | 0 | Epifaunal | Immobile |
| Arcidae | 34 | 34 | 0 | 0 | 0 | 0 | 0 | 0 | 0 | 0 | - | Epifaunal | Mobile |
| Glycymerididae | 29 | 27 | 2 | 1 | 0 | 1 | 2 | 0 | 0 | 0.14 | 0 | Infaunal | Mobile |
| Overall | 2030 | 1948 | 61 | 7 | 43 | 11 | 51 | 10 | 21 | 0.06 | 0.26 | - | - |

The relative abundance of prey family varies and is not showing any control on DF or IDF (Fig 3A-C). DF and IDF vary significantly between families (Fig 3B, C). Anomidae shows highest DF (0.17) followed by Glycymerididae, Corbulidae, Lucinidae and Cardiidae (Table 1). Lucinidae shows highest incidence of incomplete drillholes (0.7) followed by Corbulidae.

Majority of the drillholes are located in the middle (71%), followed by edge drilling (18%) and umbonal drilling (11%) (Table 1). Except for Anomidae and Glycymerididae, all the families show rarity of umbonal and edge drilling (Fig 3D).

We did not find any significant difference in DF or IDF between infauna and epifauna (Fig 4; Table 2). However, the families with incomplete drilling are all infaunal. All the complete drillholes in epifaunals are created by Naticids; Muricid drillings are observed only on infaunals (Fig 4A; Table 2). DF is significantly higher in immobile families and IDF in mobile families (Fig 4B; Table 2). Naticid drillings are significantly higher in mobile families and Muricids drillings in immobile families (Table 2).

**Fig 4.**
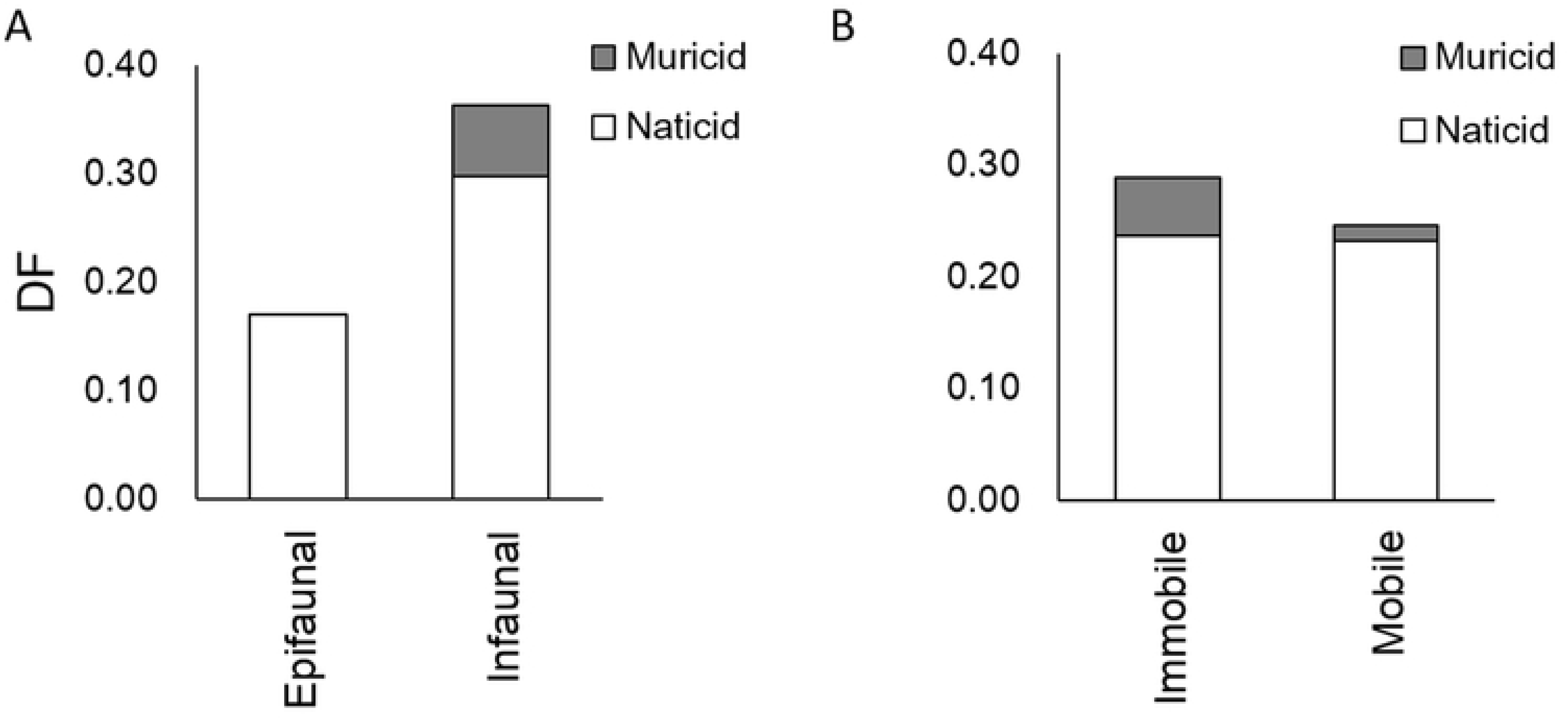
Barplots showing pooled drilling frequency (DF) for various ecological groups based on their substrate relationship (A) and mobility (B).

**Table 2.**
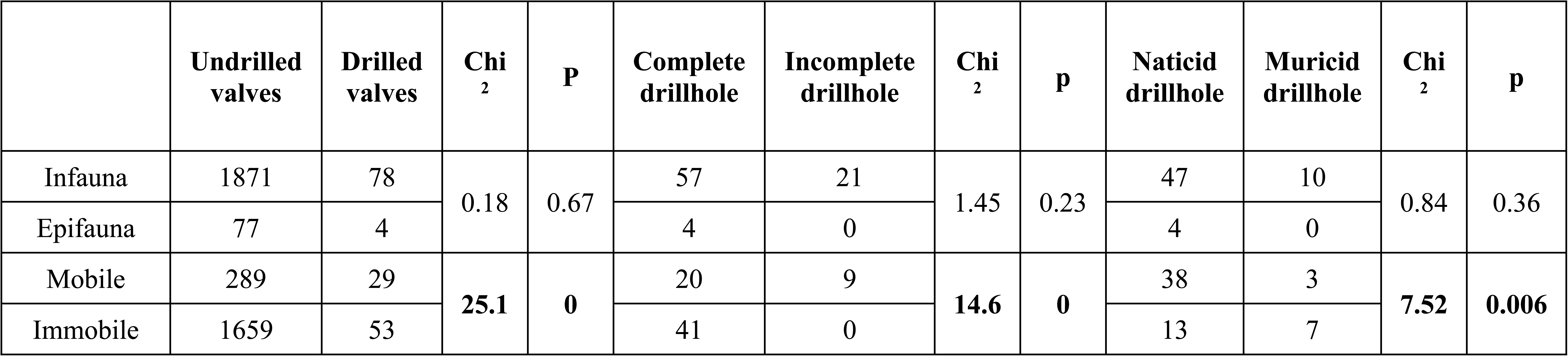
Results of Chi^2^ test to evaluate the difference in DF between various ecological groups. Significant results are in bold.

### Size selectivity

We found a significant difference in size distribution between groups with and without drilling (complete and incomplete) (Fig 5A-C; Table 3). At the family level analyses, three families (Anomidae, Cardiidae, Lucinidae) showed a significant difference in size distribution between groups with and without drilling (Fig 5D-F; Table 3). Moreover, the smaller and larger size classes showed a significantly lower incidence of drilling (Fig 6, Table 4). The three families without any drilling (Arcidae, Veneridae and Tellinidae) show significant difference in size distribution in comparison to pooled size distribution of all the drilled individuals (Fig 7A-C). Individuals are significantly larger in Arcidae (K-S test statistic D=0.57, p<0.05) than the average size of the drilled ones. Both Veneridae (K-S test statistic D=0.54, p<0.05) and Tellinidae (K-S test statistic D=0.44, p<0.05) are smaller than the drilled ones size.

**Fig 5.**
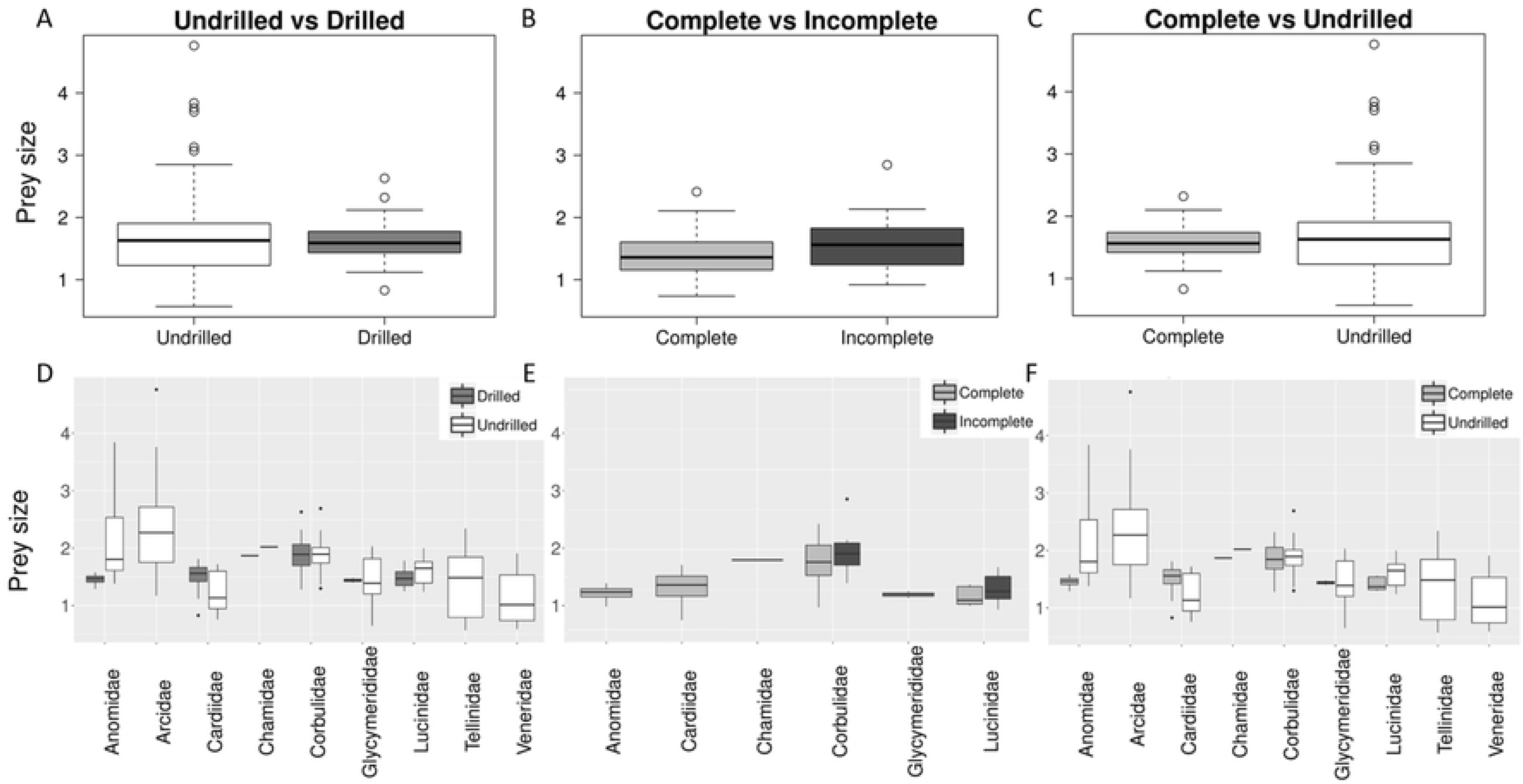
Boxplot showing comparison between prey-size of drilled (complete and incomplete) and undrilled (A); complete and incomplete (B) and undrilled and complete (C) for pooled data. Lower panel shows the same for individual families (D-F). The boxes are defined by 25th and 75th quantiles; thick line represents median value.

**Fig 6.**
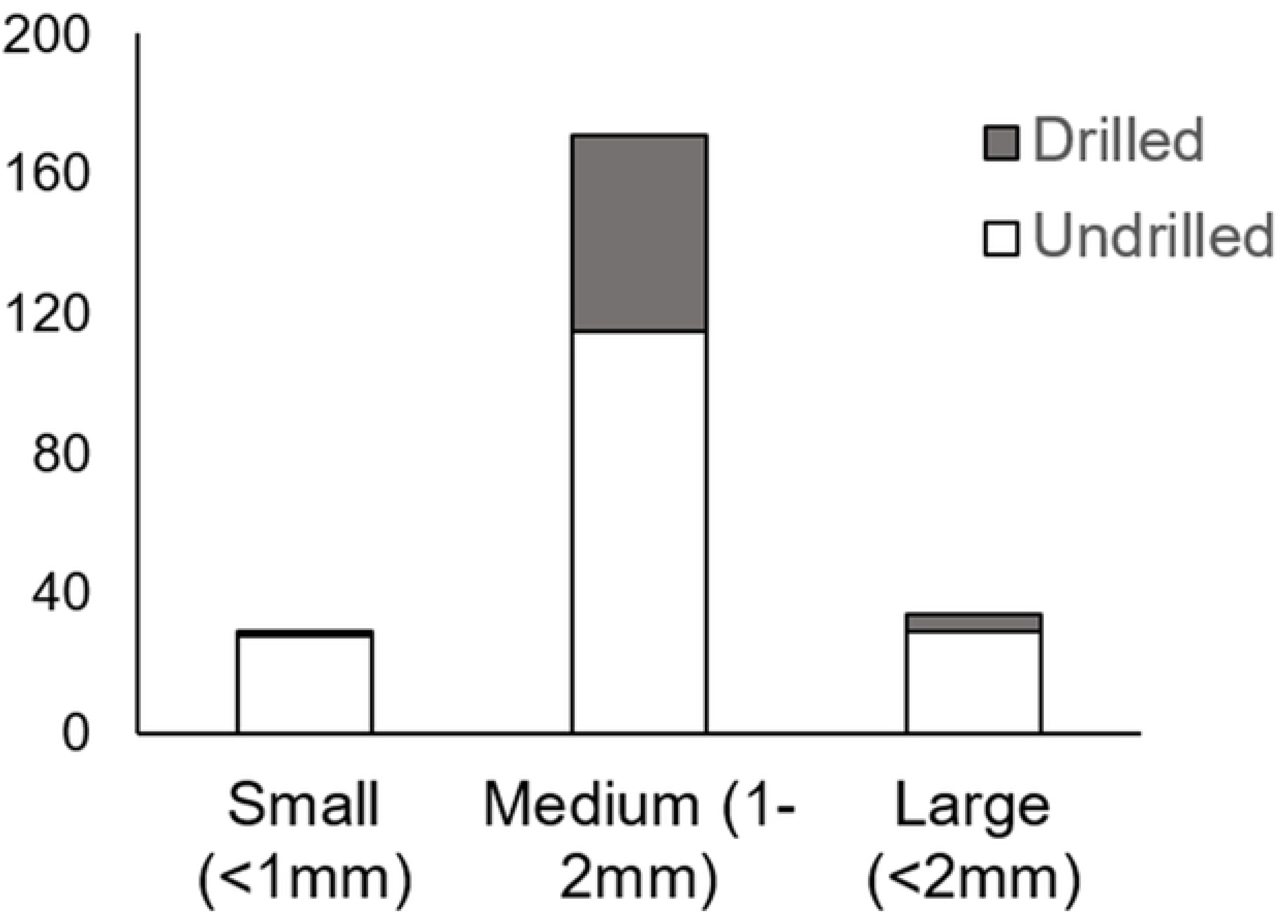
Barplot showing the proportion of drilled valves in three different size classes-small (>1mm), medium (1-2mm) and Large (<2mm).

**Fig 7.**
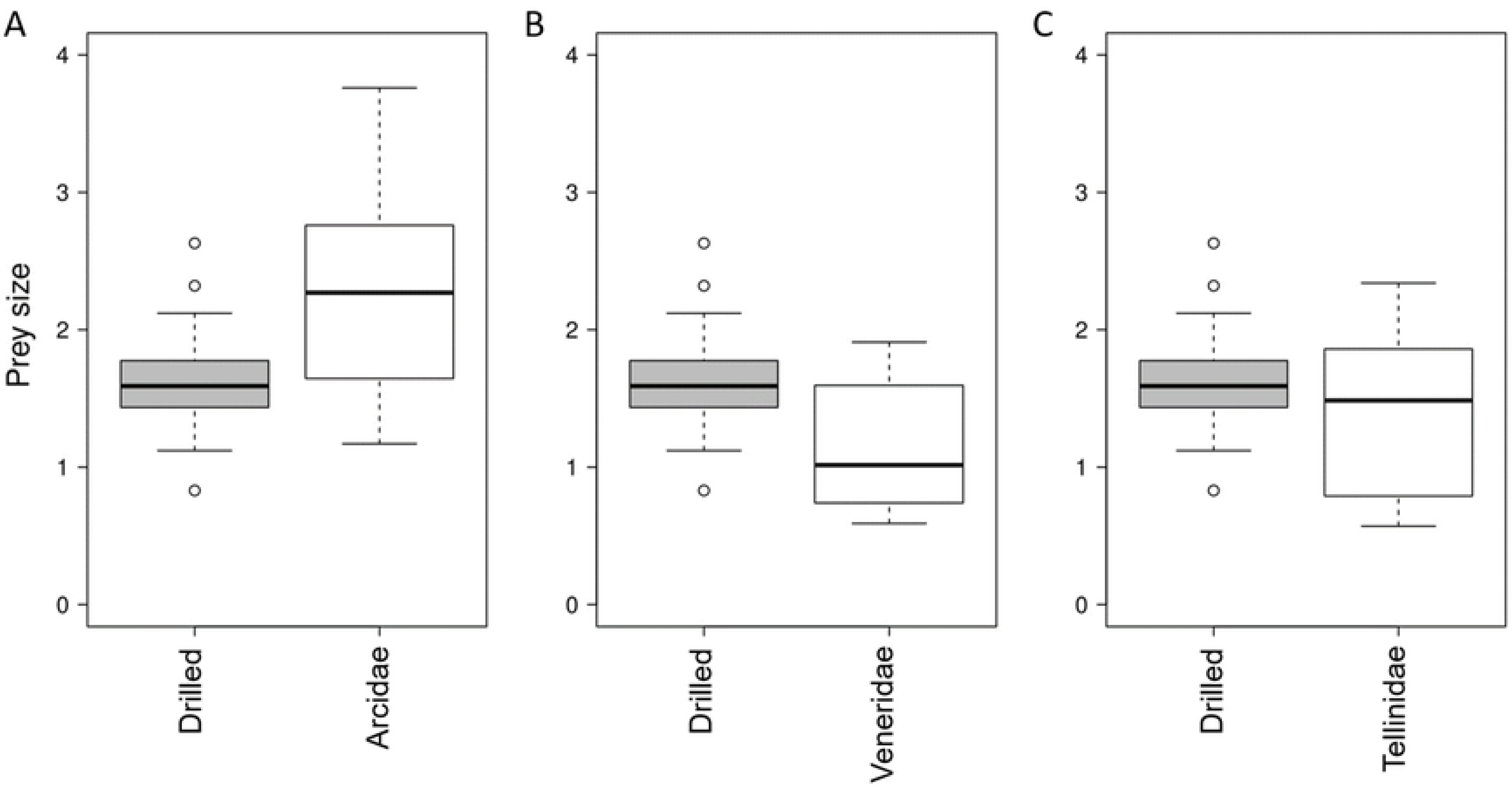
Boxplot showing comparison between pooled prey-size of drilled valves in contrast to the undrilled families inclusing Arcidae (A), Veneridae (B) and Tellinidae (C). The boxes are defined by 25th and 75th quantiles; thick line represents median value.

**Table 3.** Results of K-S test to evaluate the difference in prey size for overall and various families. Significant results are in bold.

|  | Overall |  | Anomidae |  | Cardiidae |  | Corbulidae |  | Glycymerididae |  | Lucinidae |  |
| --- | --- | --- | --- | --- | --- | --- | --- | --- | --- | --- | --- | --- |
|  | D | p | D | p | D | p | D | p | D | p | D | p |
| Drilled vs Undrilled | **0.21** | **0.02** | **0.8** | **0.03** | **0.43** | **0.02** | 0.17 | 0.9 | 0.55 | 0.64 | 0.45 | 0.05 |
| Complete vs Incomplete | 0.3 | 0.12 | - | - | - | - | 0.26 | 0.79 | - | - | 0.45 | 0.48 |
| Undrilled vs Complete | 0.2 | 0.05 | **0.8** | **0.03** | **0.43** | **0.02** | 0.21 | 0.82 | 0.55 | 0.64 | **0.7** | **0.04** |
| Undrilled vs Incomplete | 0.26 | 0.17 | - | - | - | - | 0.2 | 0.95 | - | - | 0.34 | 0.4 |

**Table 4.**
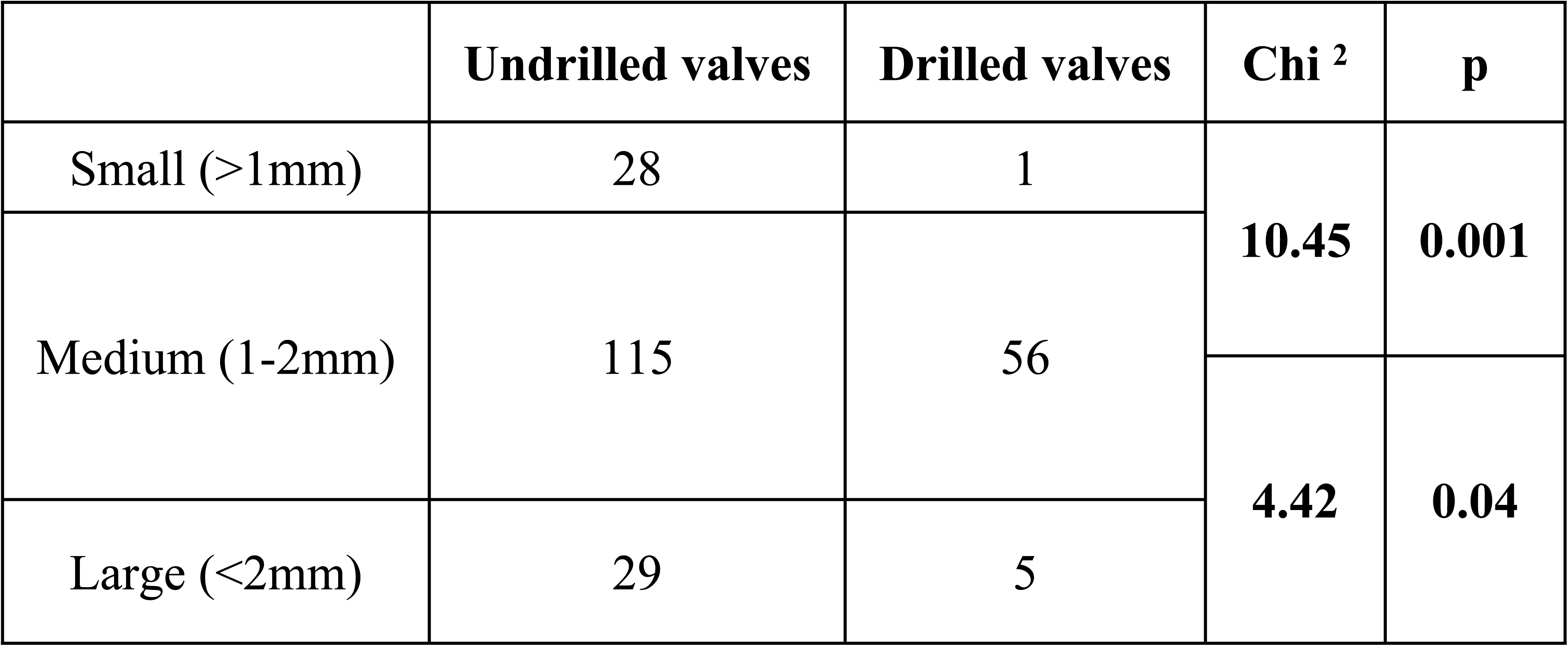
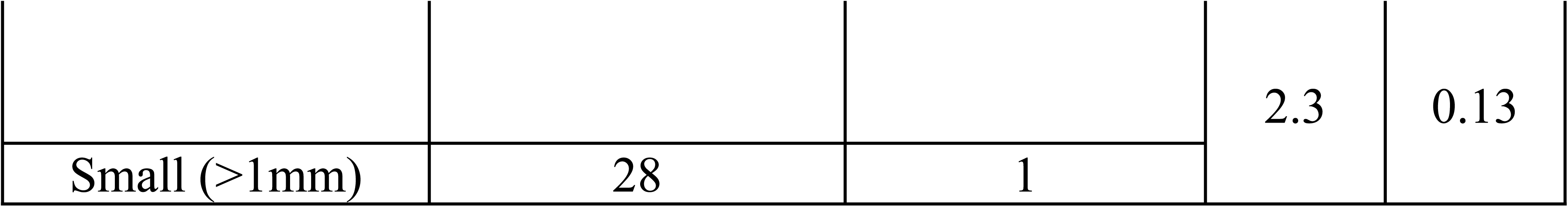
Results of Chi^2^ test to evaluate the difference in proportion of drilled valves in three different size classes. Significant results are in bold.

There is a significant positive correlation between prey size with OBD (and inferred predator size) for Naticid attacks (Fig 8A, B; Table 5), but not for Muricid attacks. However, this positive relationship does not exist for Naticid attacks on Cardiidae and Corbulidae (Fig 8C, D; Table 5). The inferred size of the Muricids is significantly larger than that of the Naticids.

**Fig 8.**
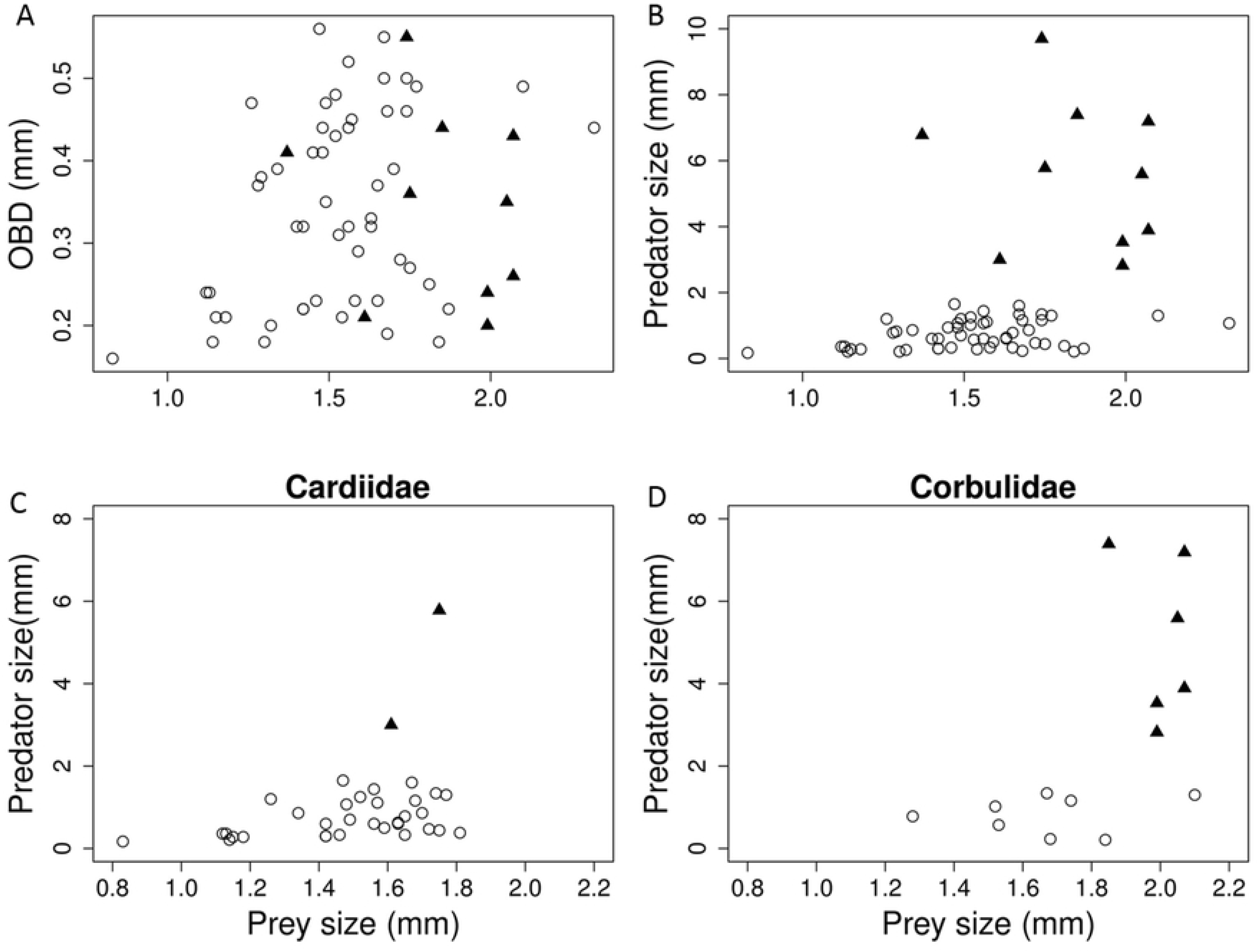
Scatterplot showing relationship between prey size and outer borehole diameter (OBD) (A) and inferred predator size (B) for pooled data. Lower panel shows the prey-predator size relationship for Cardiidae (C) and Corbulidae (D). The open circles represent Naticid attack and the solid triangles represent Muricid attacks.

**Table 5.** Results of Pearson correlation test to evaluate the relationship between predator and prey size of studied locations. Significant results are in bold.

| Location | Relationship | Pearson correlation coefficient | p |
| --- | --- | --- | --- |
| Kerala_Overall | Prey size vs OBD | **0.25** | **0.04** |
| Kerala_Overall | Prey- predator size | **0.4** | **0** |
| Kutch_Overall |  | 0.12 | 0.41 |
| Kerala_Naticid |  | **0.34** | **0.01** |
| Kerala_Muricid |  | -0.24 | 0.51 |
| Kerala_Naticid | prey/predator vs predator size | -0.85 | 0 |
| Kerala_Muricid |  | -0.93 | 0 |
| Kutch_Naticid |  | -0.57 | 0 |

### Comparison with macro fauna

The DF and IDF of the microscopic bivalves are significantly lower than those of larger bivalves of Kutch (Fig 9A, B, Table 6). The site selectivity between two provinces were also compared (Fig 9C, Table 6). In comparison to Kutch, the umbonal proportion of drillholes is significantly lower in Kerala. Unlike Kerala, Kutch fauna shows complete absence of Muricid predation.

**Fig 9.**
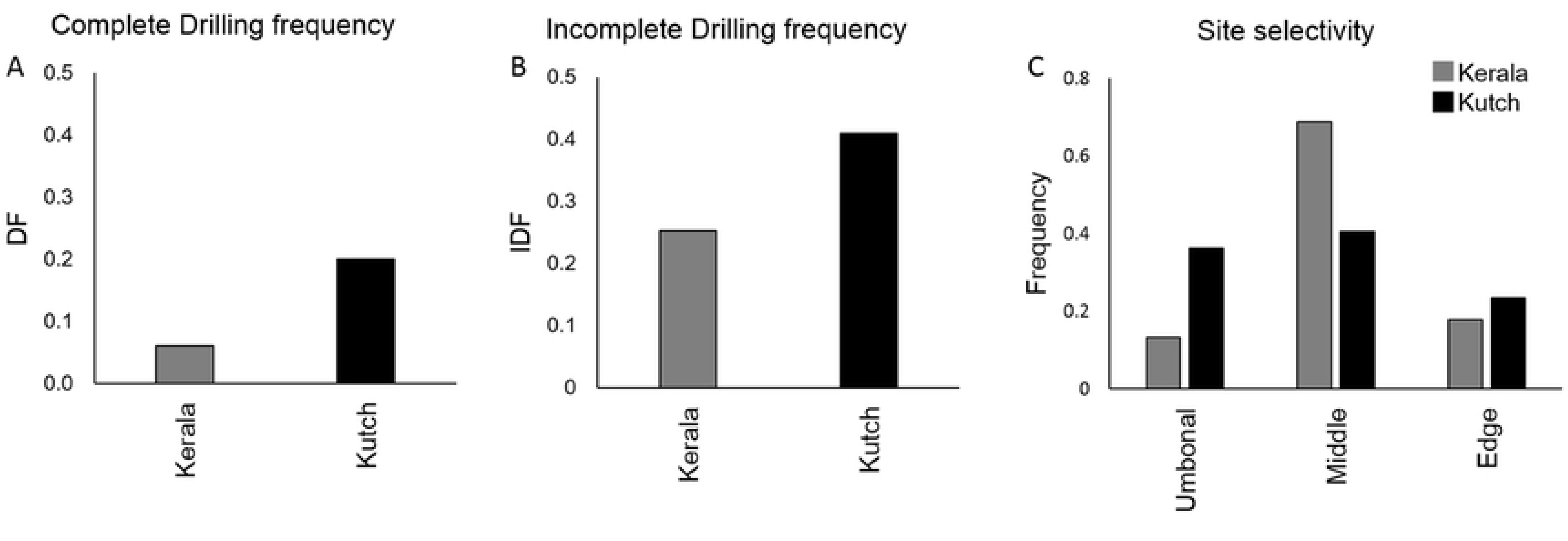
Barplots showing comparison between Kerala and Kutch drilling frequency (DF) (A); incomplete drilling frequency (IDF) (B) and site selectivity (C).

**Table 6.** Comparison of drilling frequencies between Kerala and Kutch data.

| Location | DF | IDF | Umbonal drilling | Middle drilling | Edge drilling |
| --- | --- | --- | --- | --- | --- |
| Kerala | 0.061 | 0.2 | 0.3617 | 0.4043 | 0.234 |
| Kutch | 0.253 | 0.41 | 0.1333 | 0.68889 | 0.1778 |

The two fauna show different size selectivity. Unlike microfauna of Kerala, distribution of drilled and undrilled prey size are not significantly different in Kutch fauna (K-S test statistic D=0.21, p=0.07, Fig 10A). Moreover, it does not show a significant correlation between size of prey and Naticid predator – a trend that is observed in microfauna of Kerala (Fig 10B, Table 5). Both the regions, however, show significant negative correlation between predator size with prey/predator ratio (Fig 10C, Table 5). When we compared the data with three models of increasing, decreasing and constant prey size with increasing predator size (Data File S2), both the regions matched the model corresponding to constant and increasing prey-size with increasing predator-size.

**Fig 10.**
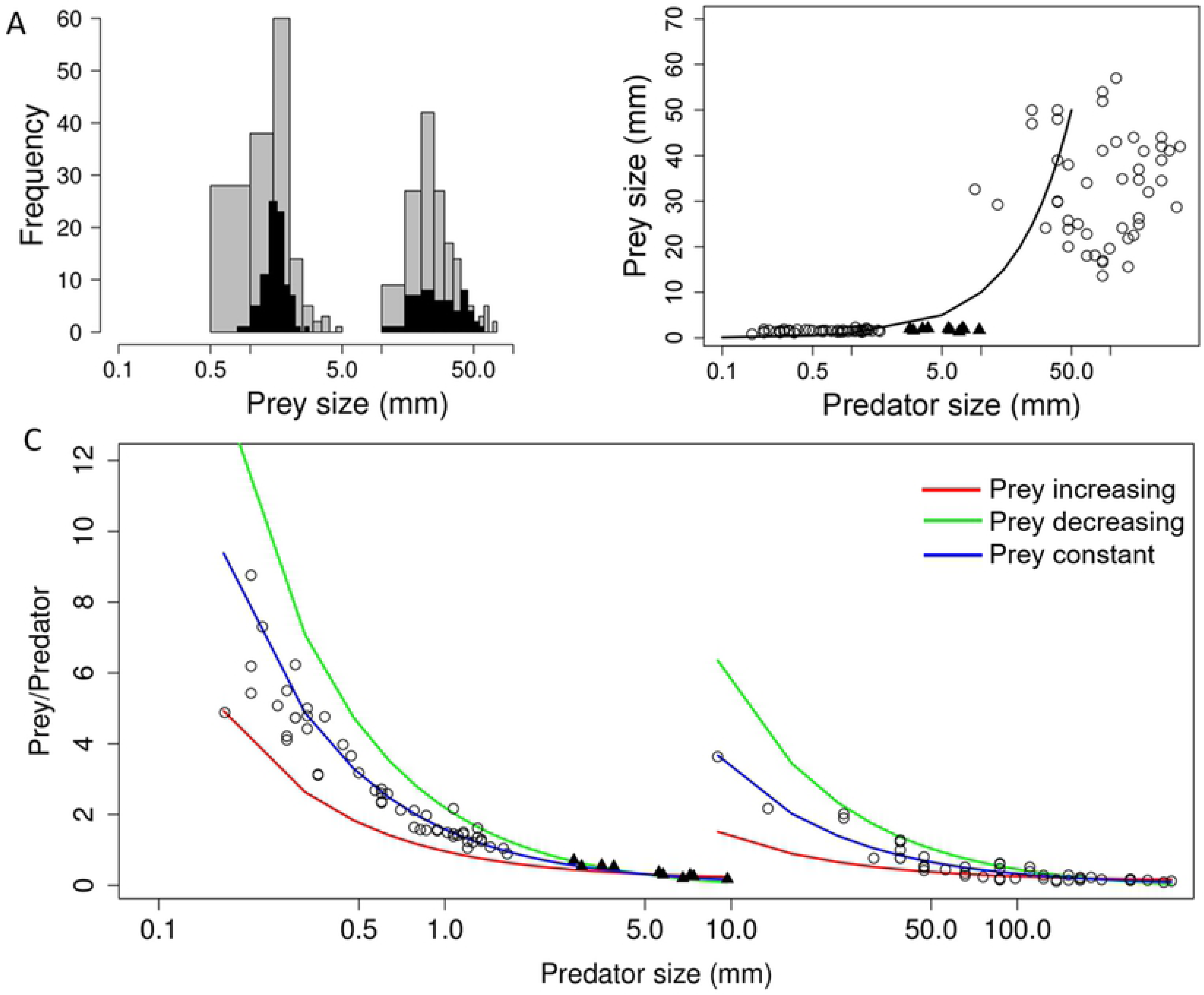
Plots showing comparison of prey size distribution between the Kerala and Kutch fauna (A), relationship between PPSRs (B) and the comparison based on the cost-benefit model (C). The open circles represent Naticid attack and the solid triangles represent Muricid attacks.

In a family specific comparison with global data of Miocene (Data File S3), microbivalves demonstrate a low DF in comparison to the mean DF in all instances except for Anomiidae; the observed DF of microbivalve is often lower than the lowest reported value of DF for corresponding family (Fig 11).

**Fig 11.**
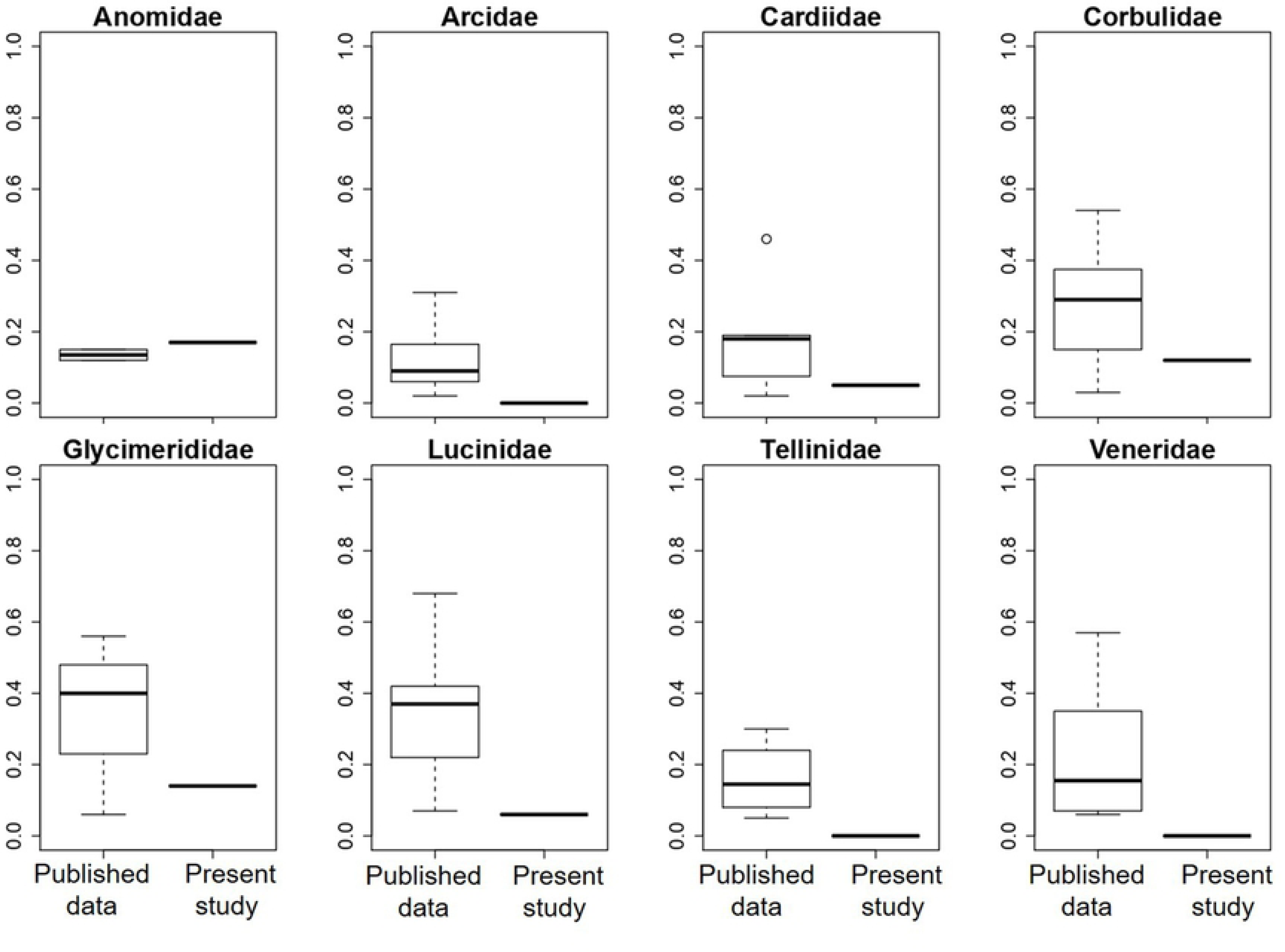
Plots showing comparison of DF for individual families between globally reported data for macrobivalve specimens (>5mm) and the present study. The boxes are defined by 25th and 75th quantiles; thick line represents median value.

## Discussion

Record of predatory drillhole is one of the unique instances where predator-prey dynamics can be studied quantitatively in deeptime. Among the studied prey taxa, bivalves are one of the major contributors [9]. Yet, documentation of predation on bivalve microfossils are largely absent except for a few brief reports [43, 44]. Consequently, the nature of drilling predation in extremely small bivalves is largely unknown. Present study is the first detailed attempt to understand the predator-prey dynamics at extreme size class.

### Nature of prey and predator

Microbivalves, primarily defined based on their small size (<10mm) [45], are enigmatic in their identity. Although some micromolluscs belong to families in which large species predominate, the majority occur in relatively few families that are composed exclusively of small species [45, 46]. On the other hand, juveniles of all families of bivalve also contribute to the microscopic size spectrum of bivalve population and they often lack adult characteristics. The studied fauna shows a number of families outside the exclusively micromolluscan families [47]. Few species found in this assemblage, are represented by larger megascopic individuals collected from the same formation [36]. The microscopic bivalves of our study, therefore, probably represent the smaller individuals of available species.

Majority of drilling in the present study is similar to those created by predatory gastropods. The exact identification of predatory family is more complicated for microbivalve community. The identification criteria for Muricid vs Naticid drillholes are primarily developed based on the morphological nature of drillings on thick shelled molluscs. With thin shelled microbivalves, the drillhole morphology is often uncharacteristic. However, many of them represent characteristic marks of muricid and naticid drilling. The existence of three genera of Naticid (*Natica, Tanea* and *Polinices*) and two genera of Muricid (*Triplex* and *Dermomurex*) from the same locality supports our identification [48]. Majority of the individuals of these predatory gastropod genera are quite small (<10mm), making them potential predators for microfauna. Experimental studies demonstrated that juveniles of both muricids and naticids create drillholes where the diameters are less than 0.1mm [49, 50], often with irregular outline [51]. Although, some of the holes in our studied material match this description (Fig 2E,G), many are quite regular in form. Moreover, a few studies claimed that gastropods are not always able to drill immediately after hatching. Muricid gastropod *Nucella lapillus* does not drill their usual prey barnacle upon hatching [52]. Instead, they feed on unfertilized eggs, small polychaetes before finally switching to their usual diet of bivalves and barnacles after attaining a minimum shell height of ∼8mm. Juvenile Naticids are also found to prey upon other groups such as ostracods [53] and foraminifera [54], instead of bivalves. Considering the dominantly smooth outline and high average value of OBD (>>0.1mm) of the studied drillholes, smaller individuals of the reported genera are more likely to be the predators over the juveniles.

Muricid gastropods attack and drill primarily epifaunally. However, they may scour shallow infaunal prey [16]. This explains the Muricid drillings on infaunal bivalves in the assemblage. Naticid is much more versatile; although primarily it hunts infaunally, it can perform the entire predatory action on the sediment surface [55]. Absolute dominance of Naticid drilling in both infauna and epifauna of the studied assemblage shows such versatility of Naticids.

### Prey selectivity in extreme size class

Leighton (2003) [56] showed a positive correlation exists between relative abundance of each species and their predation intensity for brachiopod group. However, results from other studies challenge the generality of this observation [16, 25, 41]. Our data also shows that abundance of taxa is not a good predictor of drilling frequency. We found Cardidae to be the most abundant taxa and the highest DF is observed in Anomidae indicating prey selectivity.

Incomplete drillholes are the signature of failed attacks caused by effective defensive morphology of the prey (such as large size, ornamentation), presence of secondary predators, competition etc [57–59]. Our data shows strong association between IDF and the morphological defense of the prey taxa. Incomplete drillholes occur only in two families, Lucinidae and Corbulidae, characterized by unique morphological feature. Lucinidae species have heavy ornamentation on the shell surface which is known to impart resistance against drilling [60]. Concentric ribs of Lucinids that increase effective thickness, can provide effective resistance against drilling only when selected as the drilling location. The fact that majority of the incomplete drillholes located on the ribs and not in between, supports this argument (Fig 2G). Drilling predation is claimed to be inhibited in Corbulidae due to the presence of conchiolin layers within corbulid valves [41, 61–63] along with other antipredatory morphology such as increased valve thickness, increased inflation [64]. The lower success rate in Corbulidae points to the morphological defence. We also found average prey size with incomplete drillholes are larger than those with complete ones for both of the cases; this supports the claim of bigger size as responsible factor for producing incomplete drillholes in these two families.

Ecological selectivity plays a major role in drilling predation. In general, epifauna are more susceptible to predation by epibenthic than infauna [1, 65]. Our results, however, does not support this notion of increased protection for infauna at such small size class. This may be due to the fact that depth of infauna at such size class is not enough to offer effective protection against predators including epifaunal predators. Abundance of Muricid drilling in the infaunal prey in our sample supports this. Mobile behavior of bivalves has been claimed to be an efficient strategy to evade predators [13, 66, 67]. Mobility seems to be an effective defense even at such small size class, especially fending Muricid gastropods that succeeded more attacking immobile prey. Mobile prey even at small size seems to either achieve complete evasion (low DF) or escape after being attacked (high IDF); this pattern indicates that mobility offers predation refuge for extremely small sized bivalves.

The position of the drillhole on the prey shell is indicative of the degree of behavioral stereotypy of the predator. Non-random siting of drillholes is primarily related to manipulation technique of the predator [68] and dependent on the morphology of the prey [69–71]. Such site preference is widely reported, especially for Naticids [22, 72–75]. Such high level of stereotypic behavior of Naticids may be due to the fact that they completely cover their prey inside the mantle and has lesser flexibility. Muricids, on the other hand, crawl across bivalve prey before initiation of drilling, and thus may show more variation in drillhole position [76]. We did not, however, find any difference in degree of stereotypy between Muricid and Naticid. Majority of our specimens show central position of drillhole irrespective of the predator identity. The location of the drillholes also provide important information on predation dynamics. Umbonal drilling, although common among bivalves in general, is not profitable for the predator because of the high thickness of area. This thickness increases substantially with increasing size of the prey and may pose deterrence to drilling. The common occurrence of umbonal drilling is probably due to the fact that it offers the predator some advantage in handling the prey more effectively. At extremely small size, umbonal thickness is not substantially different from the rest of the shell and hence should be preferred. Our specimens, however, do not show a significant dominance of umbonal drilling. Edge drilling, on the other hand is considered a faster technique albeit risky; a prey can damage the feeding organ (proboscis) by clamping down [77]. Edge-drilling was used when competition was intense [78]. We also found a significant number of edge drilling in our data indicating to probably a competitive scenario [12]. This is not unlikely considering the large number of young individuals after spawning season of predator that exponentially decrease with time before reaching a plate near adulthood.

### Size selectivity in extreme size class

Size is a crucial factor in controlling prey selection by drilling predators. Drillhole size is indicative of the size of the drillers for both Muricid and Naticid gastropods [22, 79]. Well accepted equations to derive predator size from OBD are primarily developed from feeding experiments using extant gastropods of normal size; the applicability of these equations to smaller size class has not been tested explicitly. The inferred gastropod size using the drillholes of our study matches the general size spectrum of gastropods reported from this locality [48] confirming the generality of these relationships even at such smaller size class.

The positive correlation between Naticid predator and prey size in microbivalve assemblage indicates a size dependent prey choice. This relationship has been explained as an energy maximizing strategy that balance the energy gain associated with the large food items and the energy spend associated with capturing and boring that large prey item [22]. For Muricid, the size relationship between prey and predator does not show any significant correlation. Unlike Naticids, energetically viable attacks by Muricids are often associated with insignificant correlation between predator and prey size [26, 57].

Previous studies reported the existence of a handling limit for a specific predator, beyond which the attacks are likely to fail and prey larger than this handing limit is immuned from successful attacks [22, 26]. Size emerges as an important factor when we evaluate the taxa without any drillholes. Both Arcidae and Veneridae are significantly different in size than the families with drillhole. The larger size of Arcidae supports the idea of a “size refuge” [1, 22]. The smaller size of Veneridae is also probably protecting it by offering limited nutritional value and hence providing a “inverse size refugia”. Although Tellinidae did not show any size difference, extremely low abundance (low encounter rate) of this taxon may be responsible for lack of drilling.

### Nature of prey-predator size dynamics

Prey preference by the predator is best evaluated using cost-benefit model developed on the tenets of optimal foraging theory. Such models have been used to correctly predict the prey choice for both Naticid [22, 41, 80] and Muricid [26]. For these models, cost-benefit ratio is estimated by the ratio of shell thickness and internal volume of the prey. Due to the small size of our specimen, it was not realistic to measure the thickness of the shell. In absence of this information, ratio of prey and predator has also been used to evaluate the relative change of predatory behavior [81]. Predators always prefer the prey item with lowest cost-benefit ratio in the size range that can be handled [16]. Cost: benefit ratio of a particular prey taxon generally decreases with ontogeny due to relatively higher energetic yield of the biomass [22]. Consequently, for a specific predator size prey with larger size ought to be beneficial if thickness does not change significantly. Because of the extremely small size and early ontogenetic stage, we do not expect significant variation in thickness in the microbivalve population. Our study, however, shows a very high prey: predator size ratio for small predators and the ratio decreases with increasing size of the predator. To give rise to this apparent counterintuitive decreasing prey-predator size ratio, the predator needs to exercise one of the following options with increasing size: a) maintain a constant prey size choice, b) choose smaller prey or c) choose larger prey, but not proportionally large. Our models show that, the declining ratio of prey:predator is achieved by maintaining a constancy of prey size or choosing inadequately large prey. Interestingly, Muricids tend to show a preference towards lower size in comparison to Naticids. In comparison to megabenthic community of Kutch, the macrobenthos is characterized by a higher prey-predator ratio although the change in prey choice by increasing size of the predator shows similar trend. The relatively lower value of prey-predator ratio for larger predators of Kutch is still probably energetically viable because energetic yield of the prey increases exponentially with increasing prey size ([22], Fig. 1)

### Possible effect of taphonomy and depositional environment

Comparison might be affected by the fact that these two regions represent two different depositional environments. Kerala has been interpreted to represent a seagrass environment [39] that is often characterized by low of predation pressure [82, 83]. The emergent seagrass blades cause decrease in the mobility of predators and the visual detection of prey [82, 84] and the roots and rhizomes of the seagrass act as a barrier to digging predators from attacking the infaunal preys [83, 85]. Thus, the predation refuge which the seagrass offers to the bivalves causes the decrease in overall predation pressure. However, the infauna in the microbivalves of the studied locality does infact show slightly higher predation intensity. Moreover, the low family-specific DF of microbivalves in comparison to global record of Miocene makes it unlikely that the depositional environment alone is creating such low predation intensity.

Taphonomic attributes are different for macro and micro invertebrates. Generally smaller individuals shelly invertebrates are poorly preserved [86] and are often affected more by other taphonomic attributes such as hydrodynamic sorting, that may even alter drilling signature [12, 87]. Considering the relatively pristine state of preservation of macrobenthos of our study, however, it is unlikely to assume taphonomy to be the primary contributor of generating the pattern.

### Evolutionary implication

In a study on modern brachiopods, Harper and Peck [33] found a significant proportion of micromorphic brachiopods in the tropics. All three extant clades of brachiopods from tropical ocean are micromorphic. Brachiopods found till 17° in the southern hemisphere are characterized by the complete absence of large (>10mm) species [88]. The micromorphic individuals were often better protected again durophagous predation. In a study on the predation intensity of temperate rhynchonelliforms, Harper et al [32] found the existence of both classical size refugia for larger individuals as well as “inverse size refugia”. Along with low frequency of repair scars in large individuals, the micromorphic ones (<5 mm) showed almost complete absence of predation. The preferred target group appeared to be the medium size class (20-40 mm) with the frequency of repair scars normally distributed around that class validating the prediction of optimal foraging theory [89]. Apart from the low energetic yield of the small brachiopods, the low intensity of durophagous predation in small brachiopods could be because of their inaccessibility due to cryptic life habit [90]. Based on these, Harper and Peck [34] hypothesized that micromorphy of brachiopods is a likely outcome of intense durophagous predation in the tropics and proposed micromorphy to be an adaptive response to increasing durophagy at lower latitudes. Our study finds evidence of effective defense rendered by micromorphy against drilling predation. The observed DF is the lowest in comparison to majority the reported DFs from Miocene [25] pointing to the rarity of such low predation intensity. Because of the effective defense in large and small groups, we can expect an evolutionary trend towards increasing variance in the size of the prey. Phanerozoic nature of predator-prey size ratio shows an increase without significant change in prey size [91]. The variance in prey size is not showing recognizable directional trend either (Fig. 2A, [90]). However, it is known that the preservation potential of small shells is significantly lower than the larger shells [86]. This makes it hard to comment on the evolutionary trend of predation in extremely small prey. Moreover, we do not know if the intensity of predation changed inhomogenously across size classes. It would be, therefore, important to focus on relative proportion of drilling frequency across size classes through time to appreciate the advantange of micromorphy against predation in evolutionary timescale.

## Conclusions

Our study documents the drilling predation dynamics in the extreme size class of micromolluscs from Early Miocene deposits of India. Our analyses demonstrate that the drilling predation in extreme size class is highly selective in terms of prey taxa, size, mobility and site selection. Drilling occur primarily on medium size class (0.83-2.32 mm) and prey outside this size range are less likely to be attacked. This indicates the existence of an “inverse size refugia” for extremely small prey along with the classical size refugia existing for large prey. Mobility is found to be deterrent to drilling predation and it also increases failure. In comparison to the predation in macrobenthos of the same biogeographic province of coeval formation, microbenthos shows a lower level of predation intensity and rate of failure. The interactions in microbenthos seems to be more strongly size-dependent compared to those among the macrobenthos that are often characterized by a lack of prey-predator size relationship. In comparison to the drilling predation in macrobenthos of global occurrences during Miocene, microbenthos shows a lower level of predation intensity in a family specific analyses. Prey-predator interaction in extreme size class highlights the importance of size in determining the nature of predation dynamics. Considering the plasticity of body size in response to environmental triggers, changing predation dynamics would be expected during times of environmental change and would have significant effect in shaping the natural selection of a group in deep time.

## Acknowledgements

This work was supported by Academic Research Grant, IISER Kolkata (ARF 2018-19), IISER Kolkata graduate research fellowship and DST Inspire fellowship.

## Supplementary materials

**Data File S1. Dataset used for present study**

Family: Family names of bivalves; Prey size: Length of the prey; Drill hole morphology: morphology of the drill holes/ Predator identity; Drill hole diameter: Outer borehole diameter of the drill hole; Position: Position of the drill hole; Completion: Completion of drill hole; Drilling: Type of prey (drilled or undrilled); Predator size: Length of the predator, Kerala; Location: Name of the location from where specimens were collected; Size class: small, medium or lage.

**Data File S2. Dataset used for present study**

Prey1_increasing (mm): Prey size in increasing order kepping minimum and maximum values same as original data; Prey1_decreasing (mm): Prey size in decreasing order kepping minimum and maximum values same as original data; Prey1_constant (mm): Mean prey size of original data; Predator1_increasing (mm): Predator size in increasing order kepping minimum and maximum values same as original data; Location: Name of the location from where specimens were collected.

**Data File S2. Dataset used for present study**

Authors: Name of the authors of the published literature from where DF is collected; Year: Year of publication; Title: Title of the literature; Prey: Type of Prey item; Prey family: Name of the famies; DF: Drilling frequency; Age: Age of the formation from where species are collected.

